# Correlative scanning electron and super-resolution structured illumination microscopy

**DOI:** 10.64898/2026.02.09.704937

**Authors:** Joseph R. Hamiliton, Summer K. Levis, Guy M. Hagen

## Abstract

Correlative microscopy techniques are used for many different applications in the biological sciences because the comparison of different imaging methods allows researchers to gain more insight and data from samples. Correlative light and electron microscopy (CLEM) methods have been developed to preserve biological samples to withstand the harsh environments necessary for electron microscopy. After first being imaged using widefield (WF) and super-resolution structured illumination fluorescence microscopy (SIM), a NanoSuit chemical treatment was applied to a mammalian testis sample before imaging with scanning electron microscopy (SEM). This was done to compare the image quality and resolution of each technique. SEM yields higher resolution and offers validation of results from SIM.

## Introduction

Scanning electron microscopy (SEM) is a method used to obtain high-resolution images of microscopic samples by coating the sample in a thin layer of conductive material and scanning the surface with an electron beam, which generates an image. A high vacuum environment is necessary for SEM but simultaneously creates undesirable conditions for biological samples by drying, dehydrating, or otherwise distorting them [1,2]. To combat these unfavorable outcomes, samples must be specially prepared prior to imaging by SEM. One such chemical treatment is the NanoSuit [3,4], which offers a layer of protection against the high vacuum environment. A modification of the original NanoSuit method, called surface shield enhancer (SSE), consists of electrolytes and glycerol and protects the sample from moisture loss and other damage from the high vacuum conditions and electron beam without compromising the quality and resolution of SEM imaging [5].

Although SEM provides information regarding the structure and surface of a sample, important functional details and insights into dynamics can only be obtained from other techniques, such as fluorescence microscopy [6]. However, optical microscopy is unable to attain the same high-resolution images that SEM does. The cooperative use of electron and optical microscopy— correlative light and electron microscopy (CLEM) allows researchers to observe and relate structural and functional characteristics of biological samples by utilizing both SEM and optical microscopy on the same region of a biological sample [7]. The SSE NanoSuit is an ideal chemical preparation for samples that undergo CLEM, as it provides the conductive and protective layer for SEM, preserves the quality of the sample, and can be removed to restore slides to their original state for further optical microscopy [8,9].

The choice of a specific optical microscopy technique is central to the success of CLEM in research applications. Fluorescence microscopy allows for 3D imaging of the inside of cells and tissues that have been tagged with fluorescent labels to gain functional information in sample but is hindered by the diffraction limit, thus generating low-resolution images. To combat this, structured illumination micrscopy (SIM) [10–14], a fluorescence micrscopy technique that uses shifting illumination patterns to generate a collection of images, was designed to accomplish the acquisition of optically sectioned and/or super-resolution images. Maximum *a posteriori* (MAP)-SIM [15] is an evolution of standard SIM methods and is useful for imagng past the diffraction limit at incresed sample depths [10]. Used to process SIM data, SIMToolbox [16] is a MATLAB software that generates widefield (WF), optically sectioned SIM (OS-SIM) [14], and super-resolution MAP-SIM images from a single set of data.

These techniques were used to image a rat epididymis tissue sample with SEM, widefield fluorescence microscopy (WF), and MAP-SIM. The high-resolution structual data from SEM and the information derived from super-resolution fluorescence imaging via SIM can be combined to enhance the strengths of each technique while minimizing the weaknesses to get as much information as possible from one sample. This can be applied to live biological samples [17–20] with use of the NanoSuit method and has applications in biomedical research, histology, pathology, and potentially numerous other research fields that utilize high resolution imaging.

## Methods

### Surface shield enhancer (SSE) NanoSuit preparation

The SSE solution was prepared by combining 5.00 g of each sucrose, fructose, and sodium chloride which were dissolved in 500 mL of deionized water. Additionally, 1.25 g of citric acid and 0.05 g of sodium glutamate were included in the solution and stirred until all solids dissolved. To create a 1:2 ratio for the final solution, 150 mL SSE was combined with 300 mL glycerol [1], the final pH was 7.4.

### Slide preparation

A few prepared slides were selected (Carolina Biological, item 31-6464, rat epididymis) and placed in toluene until the coverslip detached. The samples were rehydrated in 100, 90, and 70% ethanol, respectively, for five minutes each, and finally in deionized water for a further five minutes. The sample slides were then dipped in the SSE/glycerol solution for 10 minutes at room temperature [8]. The tissue samples were stained by the manufacturer with hematoxylin and eosin (H&E) and are brightly fluorescent.

### SEM imaging

One of the sample slides was coated with gold for 5 minutes using a sputter coating machine (Desk V, Denton Vacuum, New Jersey). Once the gold conductive layer was applied, the slide was cut (to reduce its size for the SEM chamber) with a diamond tip glass cutter, then placed into the SEM. The sample was then imaged at 30 kV using a Vega 3 SEM (Tescan, Brno, Czech Republic).

### SIM imaging

We used a home-built SIM setup based on the same design as described previously [10,11,15] (Fig. 1). The SIM system is based on a IX83 microscope (Olympus, Tokyo, Japan) equipped with a Zyla 4.2+ sCMOS camera (Andor, Belfast, Northern Ireland) under the control of IQ software (Andor). The following Olympus objectives were used: UPLSAPO 20×/0.85 NA oil immersion, and UPLSAPO 100×/1.4 NA oil immersion. The microscope is equipped with a filter set appropriate for TRITC (ET series, Chroma, Bellows Falls, VT) and a motorized X, Y, piezo Z stage (Applied Scientific Instrumentation, Eugene, OR). Figure 1 shows a simplified diagram of the SIM system.

**Figure 1.**
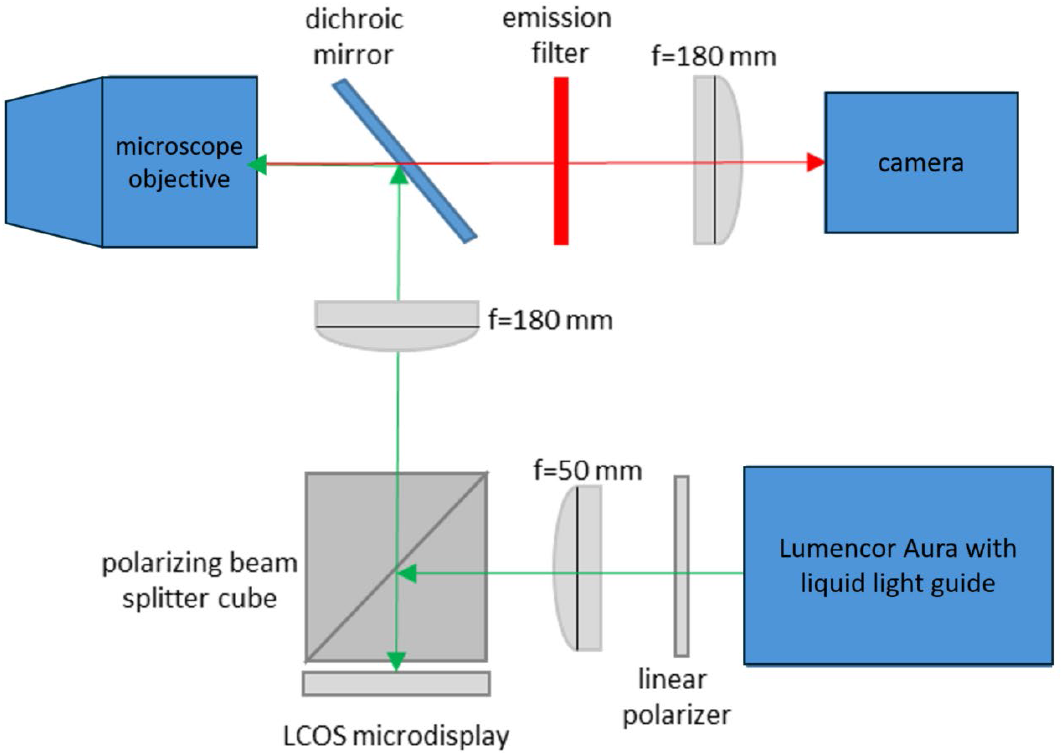
Simplified diagram of SIM system. LCOS, liquid crystal on silicon.

The SIM system uses a ferroelectric liquid crystal on silicon (LCOS) microdisplay (type SXGA-3DM, Forth Dimension Displays, Dalgety Bay, Fife, UK). This device has been used previously in SIM and fluorescence microscopy, [15,16,21–27] (among others) allowing one to produce patterns of illumination on the sample which can be manipulated by changing the image displayed on the microdisplay. The fiber-coupled light source (Lumencor Spectra-X, Beaverton, OR, USA) is switched off in between SIM patterns and during camera readout. Synchronization between the microdisplay, light source, and camera ensures rapid image acquisition, reduces artifacts, and decreases light exposure to the sample. The optical fiber was a 1.5 mm diameter multi-mode fiber (Thor Labs, Newton, NJ). The acquired SIM data was processed using SIMToolbox software [16], which generated widefield, OS-SIM, and MAP-SIM images. In some experiments (Figure 2) we used a BX53 microscope with a UPLXAPO 10×/0.40 NA objective and DP28 camera (Olympus). To evaluate the resolution of the acquired SIM images we calculated the power spectral density [28].

**Figure 2.**
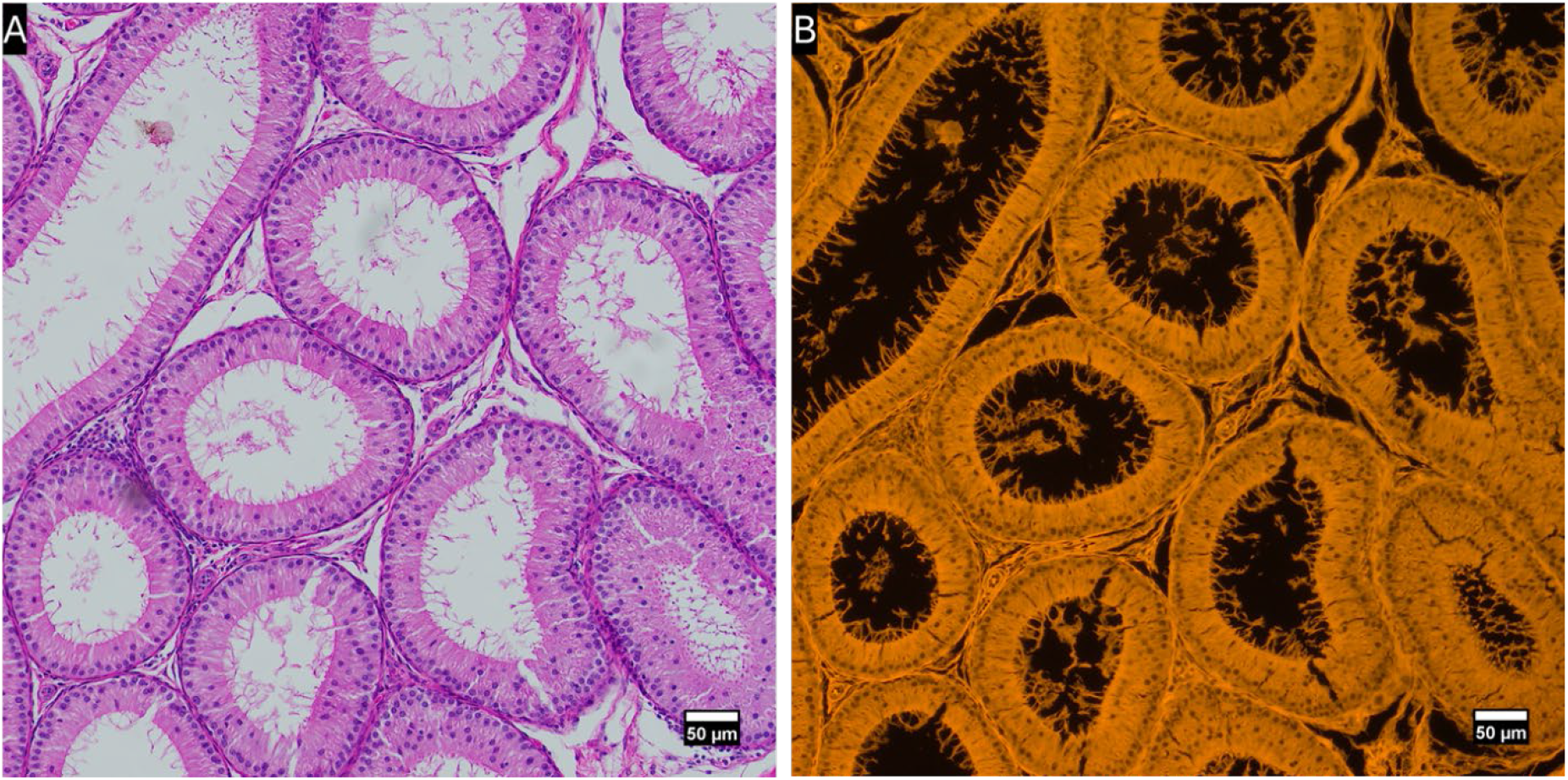
Representative transmitted light and fluorescence images of a rat testis sample using a UPLXAPO 10×/0.4 NA objective and DP28 color camera.

## Results

Both transmitted light and widefield fluorescence images of a rat testis sample stained with H&E were acquired using a DP28 color camera. This is shown in Figure 2 and offers an overview image of the type of sample used in this study.

A region of interest of a second rat testis sample was imaged with SEM (with SSE NanoSuit and gold sputtering treatment), conventional widefield fluorescence microscopy, and MAP-SIM to observe the different methods side-by-side. This is shown in Figure 3. WF and SIM imaging occurred first, followed by NanoSuit-SSE preparation and SEM.

**Fig 3:**
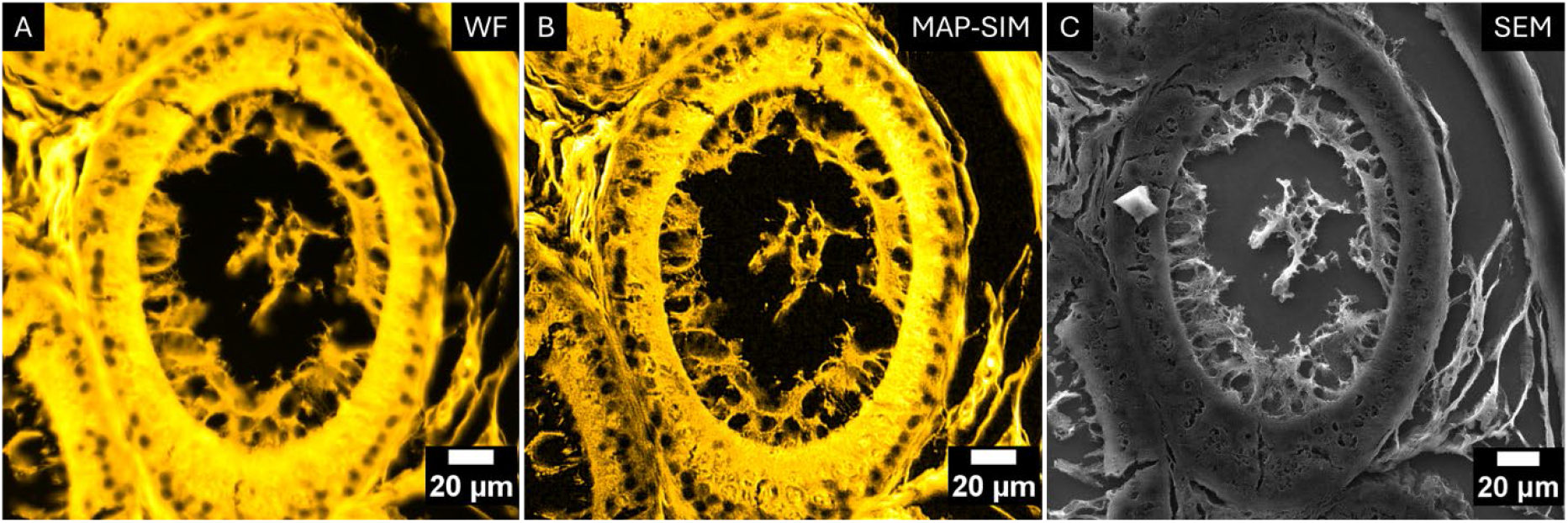
Correlative fluorescence and scanning electron microscopy of rat testis tissue stained with H&E. (A) conventional WF; (B) MAP-SIM; 20X/0.85 NA objective. (C) SEM after treatment with SSE and gold coating.

Figure 4 shows a comparison of WF and MAP-SIM acquired with a 100×/ 1.4 NA oil immersion objective and with SEM.

**Fig 4:**
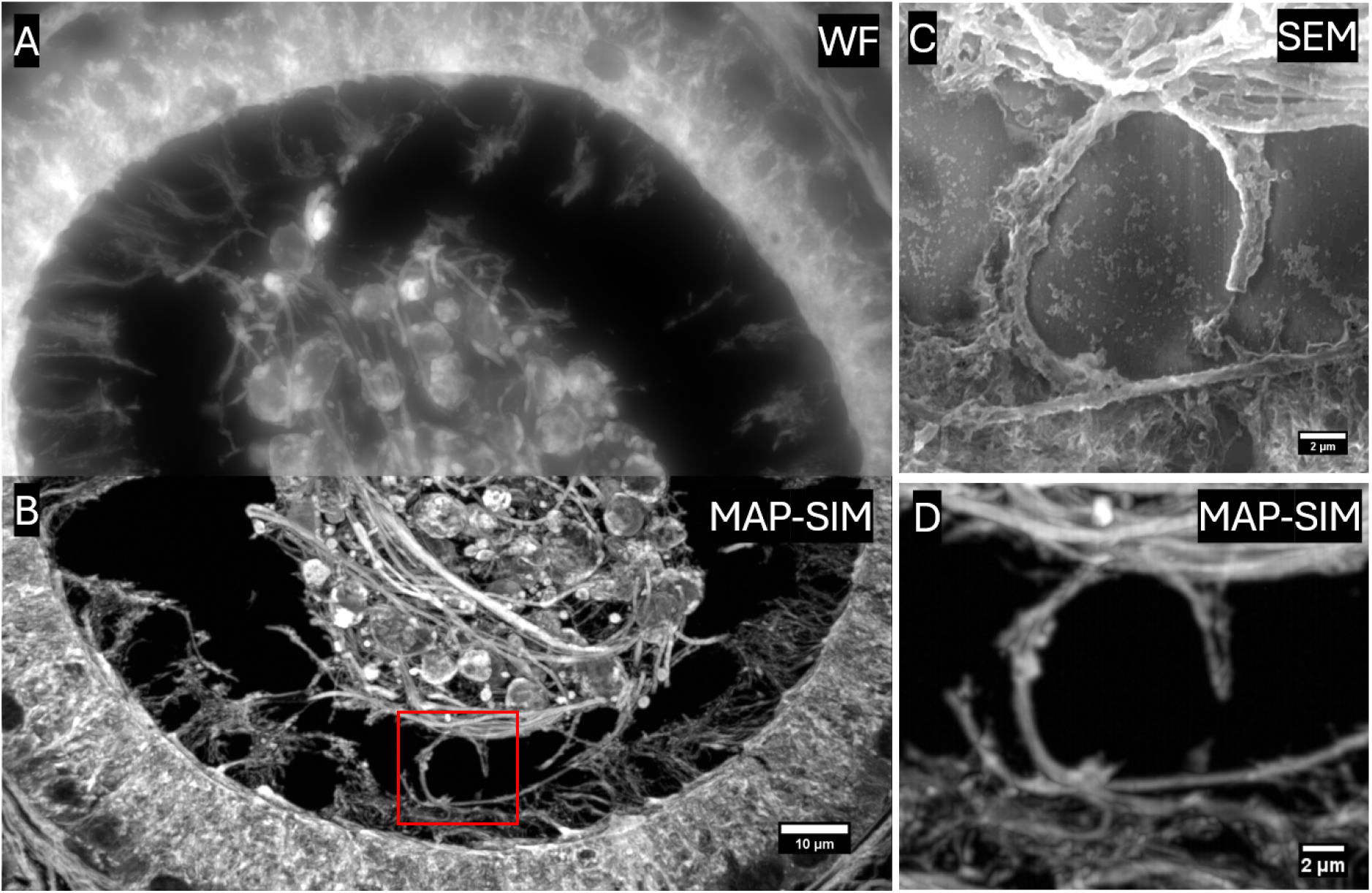
Correlative widefield, super-resolution structured illumination, and scanning electron microscopy (A) WF, (B) MAP-SIM (both 100×/1.4 NA). (C) SEM, region of interest indicated by the red box in panel B. (D) MAP-SIM of the same region of interest.

Table 1 shows resolution measurements for the images in figure 4. The resolution of each image was measured by evaluation of the power spectral density [28] as applied by us previously [10,11,29].

**Table 1:** resolution measurements of the images in Fig. 4.

|  | WF | SIM | SEM |
| --- | --- | --- | --- |
| Image pixel size | 65 nm | 65 nm | 24 nm |
| Resolution measurement | 338 nm | 142 nm | 88 nm |
| Electron beam spot size (nm) | N/A | N/A | 88 nm |

## Discussion

It has been difficult to obtain “ground truth” images when considering super-resolution microscopy. This is an important topic because both structured illumination microscopy and single molecule super-resolution microscopy (PALM/STORM type imaging [30–33]) rely on reconstruction algorithms to obtain the final image. These algorithms are under constant development, and their reliability and possible potential to introduce artifacts are of concern. Typically, conventional widefield images are shown in comparison to super-resolution images that were obtained by whichever method is being developed or applied. However, comparison of super-resolution microscopy images with SEM provides a higher resolution image for comparison and helps prove that the super-resolution optical microscopy image is accurate and free from artifacts. The results shown here indicate that super-resolution structured illumination microscopy, as implemented by us (MAP-SIM), is showing the correct sample structure and is free from apparent artifacts. Other SIM methods are more prone to pattern-related artifacts [34].

Other correlative microscopy techniques differ in procedure but serve a similar function in obtaining information about a sample from various methods. Another type of CLEM involves the use of laser scanning confocal microscopy (LSCM) followed by transmission electron microscopy (TEM) [35]. The images captured from TEM and LSCM can be overlaid into an integrated image [36], providing, for example, ultrastructural imaging of cells rapidly frozen at a moment in time at which a particular cellular function is taking place.

## Conclusion

Procedures for correlative microscopy techniques have been in development for many years and are constantly being altered to maximize the benefits from their use. The utilization of super-resolution SIM with SEM is an especially promising form of CLEM due to the relative ease of the method and surprisingly good results as shown here.

## Abbreviations

H&E: hematoxylin and eosin
MAP-SIM: maximum *a posteriori* probability SIM
SSE: surface shield enhancer
NA: numerical aperture
LCOS: liquid crystal on silicon
CLEM: correlative light and electron microscopy
SEM: scanning electron microscopy
SIM: structured illumination microscopy
WF: wide field.

